# Multidimensional Boolean Patterns in Multi-omics Data

**DOI:** 10.1101/2021.01.12.426358

**Authors:** George Golovko, Victor Reyes, Iryna Pinchuk, Yuriy Fofanov

**Affiliations:** UTMB; PennState Health

**Keywords:** Microbiome, Multidimensional Boolean patterns, Multiomics data, Co-exclusion, Co-presence

## Abstract

**Motivation:** Virtually all biological systems are governed by a set of complex relations between their components. Identification of relations within biological systems involves a rigorous search for patterns among variables/parameters. Two-dimensional (involving two variables) patterns are identified using correlation, covariation, and mutual information approaches. However, these approaches are not suited to identify more complicated multidimensional relations, which simultaneously include 3, 4, and more variables.

**Results:** We present a novel pattern-specific method to quantify the strength and estimate the statistical significance of multidimensional Boolean patterns in multiomics data. In contrast with dimensionality reduction and AI solutions, patterns identified by the proposed approach may provide a better background for meaningful mechanistic interpretation of the biological processes. Our preliminary analysis suggests that multidimensional patterns may dominate the landscape of multi-omics data, which is not surprising because complex interactions between components of biological systems are unlikely to be reduced to simple pairwise interactions.

## Introduction

Virtually all biological systems are governed by complex relations between its components. Identifying such relations involves a rigorous or heuristics-based search for *patterns* among *variables/parameters* of a system. Several algorithms have been developed to identify two-dimensional (involving two *variables*) *patterns* using correlation, covariation, and mutual information approaches. However, complex biological systems may have much more complicated multidimensional relations, which can only be described using patterns that simultaneously include 3, 4, and more variables. The main challenges in the search for such *multidimensional patterns* include computational complexity of the search; discrimination between statistically significant patterns from those which may be observed in large data sets simply by chance; and integration of heterogeneous data types (integer, Boolean, categorical, etc.) in a single pattern. This manuscript presents an attempt to address some of these challenges using *multidimensional Boolean patterns* and novel algorithms to perform the search for all possible 2, 3, and 4-dimensional patterns in multi-omics data.

## Results

### Multidimensional Boolean patterns

Boolean patterns are widely used to describe relations between components (*features*) of complex biological systems, especially when such a relation cannot be directly represented by scores based on values of variables such as by correlation, covariance, or co-exclusion ^1^. For example, in environmental microbial communities (MC), the functions vital to all microorganisms (e.g., nitrogen fixation) are usually performed by a single species. As a result, the pattern describing relations between these species and amy other members of MC cannot be observed as a correlation/covariation but is rather expected to be manifested as Boolean *one-way relation* patterns ^1,2^, where the presence of the “dependent” variable(s) requires the presence of another “provider” variable, but not vice versa (Figure 1c). Similarly, other Boolean pairwise relations, such as *co-presence* and *co-exclusion,* may be represented as Boolean patterns (Figure 1a, b). In fact, out of 16 ( ) possible combinations of the presence/absence profiles between two variables, only four may be interpreted as possible relations: *co-presence*, *co-exclusion*, and two *one-way relations* (variable 1 needs variable 2 to be present and the opposite: variable 2 needs variable 1 to be present).

**Figure 1.**
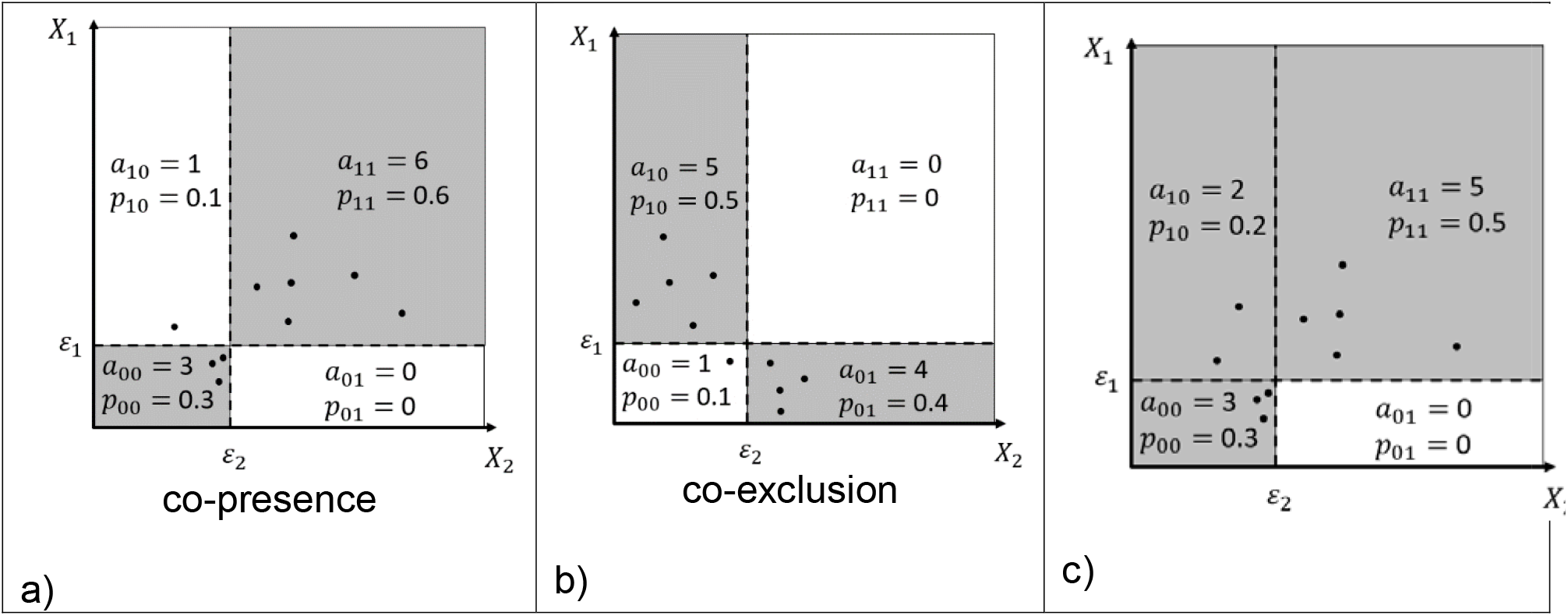
Examples of Boolean two-dimensional patterns: (a) *co-presence*; (b) *co-exclusion*; and (c) *one-way relations*. and are abundances of features (variables). *X*_1_ and *X*_2_ are the abundances of features; *ε*_1_ and *ε*_2_ represent the presence/absence threshold; *p*_00_, *p*_01_, *p*_10_, and *p*_11_ are the proportion of points (observation) located in each partition. Grey color and pattern description represent partitions requiring the proportion of observation to exceed the minimum threshold and areas (partitions) contributing to the pattern score.

In general terms, the Boolean pattern ( ) can be defined as a presence/absence (0/1) profile of Boolean variables associated with each combination of indexes ( ), so for two variables, the 2D pattern can be defined as:

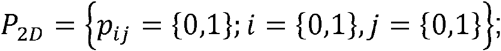

For example, the co-presence patterns (Figure 1a) is defined by the Boolean presence (1) or absence (0) values in all four partitions of the 2D space:

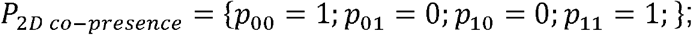

Similarly, for higher dimensional patterns (3D, 4D, etc.):

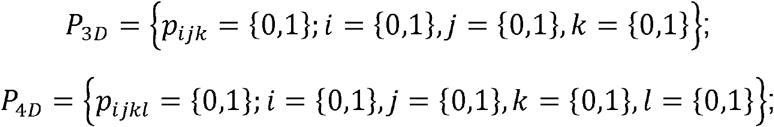

For example, 3D co-presence and two different types of co-exclusion patterns (Figure 2a,b,c) can be defined as:

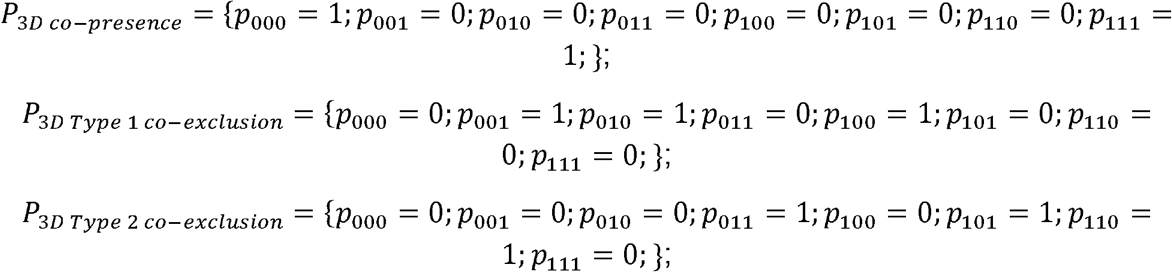

**Figure 2.**
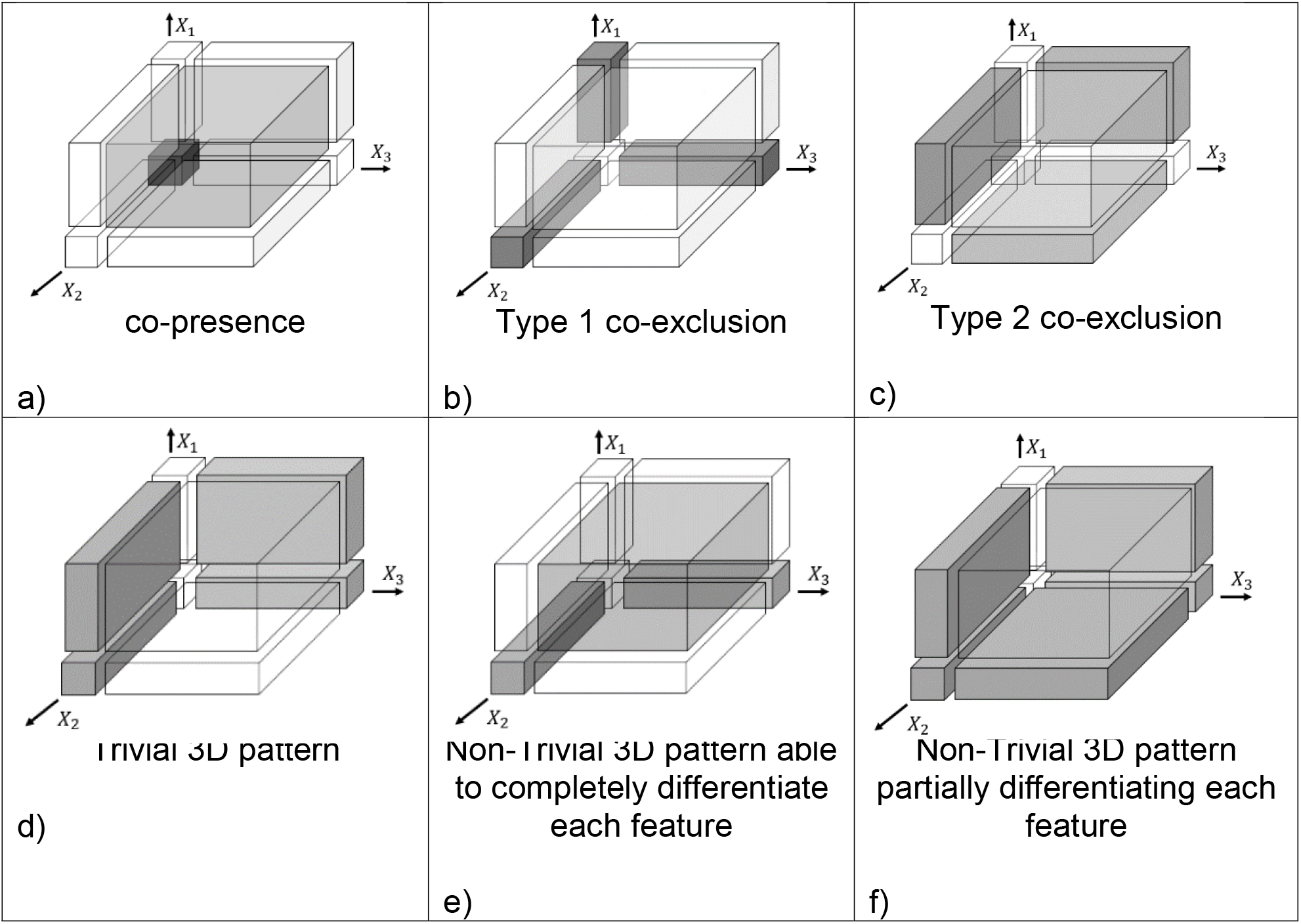
Examples of Boolean three-dimensional patterns. (a) co-presence, (b) Type 1 co-exclusion, (c) Type 2 co-exclusion, (d) Trivial 3D pattern, (e) Non-Trivial 3D pattern able completely differentiate each feature, (f) Non-Trivial 3D pattern partially differentiating each feature. Grey color and pattern description represent partitions requiring the proportion of observation to exceed the minimum threshold and areas contributing to the pattern score

### Patterns - SubPatterns Consistency

Each *n*-dimensional pattern can be decomposed into a combination of 2*n* patterns of lower (*n* − 1) dimensions. Such subpatterns can be described as a “slice” of an *n* dimensional pattern for a specific value of one of the variables. For example, each 3D pattern

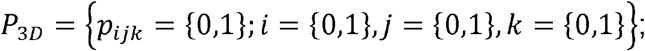

contains six 2D subpatterns:

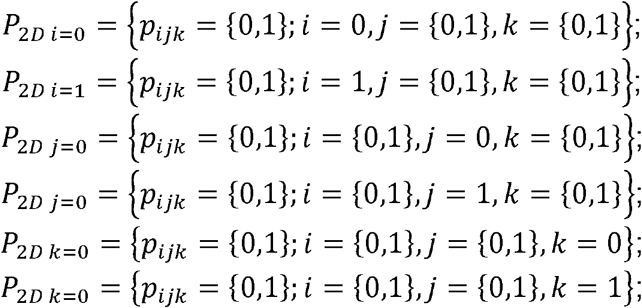

It is important to emphasize, however, that not every combination of dimensional patterns expressed between variables are possible. Each combination of dimensional patterns leads to three possible outcomes:

1. The combination of dimensional patterns leads to a unique -dimensional pattern;
2. The combination of dimensional patterns leads to several -dimensional patterns;
3. The combination of dimensional patterns leads to no -dimensional pattern, thereby being inconsistent.

For example, the following set of 2D patterns is inconsistent (cannon exist) because the corresponding 3D pattern is impossible.

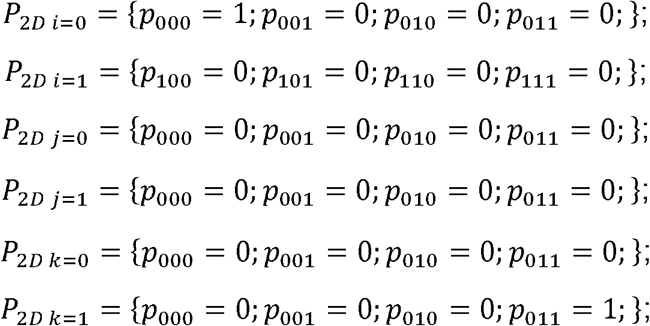

Such a relation between lower and higher dimensional patterns offers an interesting opportunity to evaluate the overall consistency of the set of patterns describing each given biological system and allows detection of contradictions between patterns. Additionally, the presence of higher dimensional patterns could help to identify the presence of missed lower-dimensional subpattern (*s*).

It is also important to mention the special role of the *p*_00_, *p*_000_, *p*_0000_ etc. partitions in the pattern definition. In some cases, such as abundance of particular microorganisms in a microbial community, the observations in this partition can be interpreted as irrelevant (both features/values under consideration are absent); on the other hand, for features such as SNPs, or proteins, however, the absence of the features has to be considered as an important part of the pattern.

### Trivial vs. non-trivial patterns

While the total number of *n*- dimensional patterns is 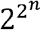, it is important to keep in mind that not all patterns can be interpreted as a possible relation between variables. For example, the 2D pattern in which *p*_*ij*_ = 1 for all combinations of *i, j* (all partitions are present) is *trivial* and can not be interpreted as any meaningful relations. Two simple criteria can be used to exclude such trivial (not allowing any relation-like interpretation) patterns:

1. For a pattern to be *non-trivial,* each variable in it must be observed in both presence (1) and absence (0) states. For example, in the case of a 2D pattern:

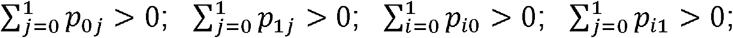 In other words, if a variable is only present or only absent in all observations, it cannot be part of any non-trivial pattern.
2. For a pattern to be *non-trivial, for* each variable its (*n*−1) dimensional subpattern when this variable is present (1) must be different from the (*n*−1) dimensional subpattern when this variable is absent (0). For example, 3D pattern:

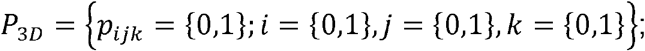

containing six 2D subpatterns:

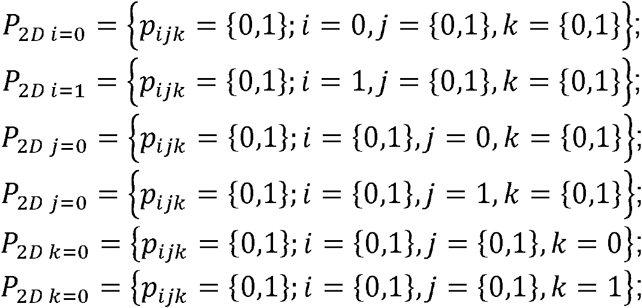

will be considered non-trivial if: *P*_2*D i*=0_ ≠ *P*_2*D i*=1_ and *P*_2*D i*=0_ ≠ *P*_2*D i*=1_ and *P*_2*D i*_=0 ≠ *P*_2*D i*=1_ simultaneously. Example of trivial and non-trivial 3D patterns can be found in Figure 2: the pattern in Figure 2e is non-trivial, while the pattern in Figure 2d is trivial because 2D subpatterns are the same for both values of *X*_1_.

One special type of multidimensional pattern-subpattern relations occurs when the *n*-dimensional pattern is a direct consequence of its subpatterns. For example, a 3D type 1 co-exclusion pattern (figure 2b) may be considered a result of three 2D co-exclusions. Such patterns be can be easily detected and excluded from consideration by requiring n-dimensional pattern to have a unique (only one) way *n* − 1 dimensional pattern representation. However, in the presented study, we decided not to exclude such patterns and keep all the "meaningful" patterns identified in real data.

### Patterns which can distinguish selected features

The analysis of biomedical data is often focused on the identification of patterns allowing to discriminate (wholly or partially) between the presence/absence of selected features. One can search for a combination of features that can identify a specific phenotype. For example, the composition of several SNPs which can be predictive genetic markers of cancer; patterns in gut microbial composition associated with Crohn disease; or combinations of environmental and genetic factors associated with the development of autism. To identify if a given *n*-dimensional pattern can discriminate feature of interest, one can perform a comparison between two (*n* − 1)-- dimensional subpatterns appearing for each value (0/1) of that feature. For example, the 2-dimensional pattern in Figure 1a can use the value of one feature (*X*_1_) to “predict” the value of another (*X*_2_). In contrast, some patterns can be used to discriminate selected features only partially/conditionally. Figure 1b shows an example where the presence of one feature (*X*_1_ = 1) cannot be used discriminate *X*_2_: it can be either 0 or 1; but when *X*_1_ =0 the only possible value for *X*_2_ is 0. There are two types of partially discrimination patterns: patterns conditionally discriminating the presence (1) or absence (0) of the feature.

The following criteria/definition can be formulated to determine if a given pattern can completely differentiate the feature.

*Feature (X*_1_) *can be completely differentiated by pattern P*_*ijk...*_ *if and only if p*_0*jk...*_ + *p*_1*jk...*_ ≤ 1 *for every combination of j, k, etc.* In other words, to completely discriminate the feature, the corresponding presence/absence values of the subpatterns appearing for each value (0/1) of this feature must never be co-present.

Partial feature discrimination can be defined by relaxing complete differentiation criteria:

a. *To be able to conditionally (partially) discriminate the presence of the feature, corresponding subpatterns (P*_0*jk...*_ *and P*_1*jk...*_) *must have at least one partition (the combination of j,k,...) where corresponding values p*_0*jk...*_ = 0 *and p*_1*jk...*_ = 1 *simultaneously.*
b. *To be able to conditionally (partially) discriminate the absence of the feature, corresponding subpatterns (P*_0*jk...*_ *and P*_1*jk...*_) *must have at least one partition (the combination of j,k,...) where corresponding values *p*_0*jk...*_ = 1 *and p*_1*jk...*_ = 0 simultaneously.*

Figure 2e shows an example of the pattern that completely differentiates (discriminates) each feature. In contrast, the pattern in Figure 2f presents an example of complex partial discrimination: it can predict the absence of *X*_1_ if *X*_2_ and *X*_3_ are present; the presence of *X*_3_ if *X*_1_ and *X*_2_ are absent; the presence of *X*_3_ if *X*_1_ present *X*_2_ are absent; absence of *X*_3_ if *X*_1_ and *X*_2_ are present; the presence of *X*_2_ if *X*_1_ and *X*_3_ are absent; the presence of *X*_2_ if *X*_1_ is present and *X*_3_ is absent; and absence of *X*_2_ if *X*_1_ and *X*_3_ are present.

Table 1 present the summary of the discriminative properties of 2, 3, and 4-dimensional patterns. The complete list of all patterns and their characteristics is available in the Supplementary Table 1.

**Table 1.** The number of different types of patterns in 2, 3, and 4 dimensions.

| Pattern Type | 2D | 3D | 4D |
| --- | --- | --- | --- |
| Total number of patterns | 16 | 256 | 65,536 |
| Non-trivial patterns p0.. included | 6 | 174 | 62,224 |
| Non-trivial patterns p0.. excluded | 4 | 96 | 31,479 |
| Non-trivial patterns able to differentiate single feature (p0.. included) | 2 | 38 | 5,401 |
| Non-trivial patterns able to differentiate single feature (p0.. excluded) | 1 | 15 | 1,909 |
| Non-trivial patterns able to conditionally differentiate the feature's presence (p0.. included) | 4 | 136 | 56,823 |
| Non-trivial patterns able to conditionally differentiate the feature's presence (p0.. excluded) | 2 | 66 | 27,770 |
| Non-trivial patterns able to conditionally differentiate feature's absence (p0.. included) | 4 | 136 | 56,570 |
| Non-trivial patterns able to conditionally differentiate feature's absence (p0.. excluded) | 3 | 81 | 29,522 |

### Pattern strength (Score)

In the search for Boolean patterns in experimental data, the pattern strength can be defined based on the fraction of observations falling in each “present” (*p*_*ij ...*_ = 1) partition of the given pattern. For example, for 2D patterns, the pattern strength (score) will depend on the fraction of the experimental observations in four partitions of two-dimensional space: *a*_00_, *a*_01_, *a*_01_, *a*_11_ (Figure 1a-c).

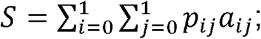

In other words, the pattern score can be defined as the fraction of observations belonging to the pattern.

Several approaches can be used to transform abundance values to the presence/absence profile ^3^. We, however, believe that since the same feature may be involved in multiple processes, each of which may require it to present in different abundance, the identification of the presence/absence threshold has to be pattern and features combination specific and require threshold (*ε*_*i*_) optimization:

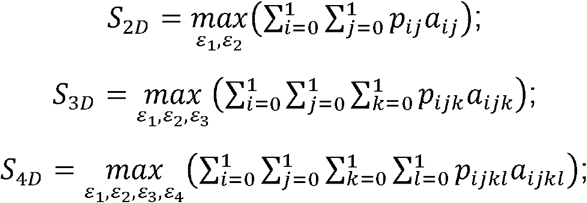

While the threshold optimization allows for embracing the possibility that different variables may have different and even pattern-specific thresholds, it can also produce some misleading results if one allows partitions required to be present in the pattern to be empty or contain very small number of observation. This effect can be eliminated by requiring a proportion of the observations (*a*_*ij...*_) in all quadrants present (*p*_*ij...*_ = 1) in the pattern to be present above a predefined *minimal presence threshold (d)*.

### Statistical Significance and Type 1 Error

For Boolean patterns, number of samples and features, choice of the minimal presence threshold (*m*), and minimal pattern strength/score (*S*_*min*_) required can significantly affect the results of the analysis by altering the number of detected patterns as well as the statistical significance of identified patterns (e.g., Type 1 error). In the presented below analysis of gene composition and microbial abundance data, several steps have been performed to minimize the probability of the appearance of such random patterns in the resulted analysis. For each pattern under consideration, instead of making an arbitrary choice of the *minimal presence threshold* (*d*) and *minimal pattern score required* (*S*_*min*_), we performed several cycles (repetitions) of the search for each pattern in a randomized (shuffled) dataset using different combinations of *threshold* (*d*) and *minimal score* (*S*_*min*_).

Counting the number of patterns passing both thresholds in randomized data allows to identify (*S*_*min*_,*d*_*max*_) combinations for which the total number of random patterns is below the predefined minimum (in the presented cases, it was set to zero) and use only these combinations of thresholds in the search for a given pattern in experimental data. It is necessary to mention that the shuffling method must reflect the underlying assumption about what is considered as the random alternative to the observed dataset (Zero Model) ^4, 5^. In the presented cases, to accommodate the fact that some features may be represented by values always lower or always higher than average, the shuffling must be performed across individual feature profiles. The analysis of the identified patterns’ statistical significance is performed by applying bootstrapping across values of individual features for all identified patterns and estimating the pattern’s p-value (by counting fraction of times when the randomized data simultaneously produces better thresholds and score then original data).

### Patterns search and optimization

The critical challenge of the search for 3, 4, and higher-dimensional patterns is its high computational complexity. Depending on the implementation, the search for each pattern could reach the complexity of *O*(*f*^*n*^*m*^*n*^), where *f* is the number of features, *m* the number of samples and *n* is the pattern dimension. Even with basic parallelization by calculating each model for each combination of features simultaneously, the naïve and straight forward implementation of the proposed approach will require an enormous amount of computational power.

In the in-house software used in the analysis presented below, significant effort was involved in improving the performance of the application. It includes early identification and filtration of the features which will not be able to fit a given model, precomputing of the feature-specific optimization grid steps, early exit from bootstrapping, and the analysis of randomized datasets. All the examples presented in this manuscript have been calculated using a single processor workstation, where analysis of all 2D patterns takes 2-4 hours and 3D patterns 1-3 days.

To demonstrate the presence of 2 and 3-dimensional patterns in experimental data, we selected one dataset where the features have been represented by numerical (stool microbiome composition) and discrete presence/absence (*H. pylori* gene composition) values.

### 3D patterns in stool microbiome composition

The stool microbial community compositions used for this analysis originated from the NIH Human Microbiome Project. Microbial profiles have been downloaded from the project website (HMQCP–QIIME community Profiling v13 OTU table) ^6^. Samples with significantly low (less than 2,000) and high (over 50,000) number of sequencing reads were excluded from the analysis. The microbial profiles of the remaining samples have been normalized against the total number of reads in each sample and transformed into relative abundance profiles merged to genus taxonomy level. Genera present in less than 5% of samples have been excluded from the analysis. No 2D patterns have been identified in the data with a required pattern score of 0.99 and a minimum presence threshold of 0.15. A total of 195 3D patterns have been found (Supplementary Table 2). Figure 3 shows 3D patterns in stool Human Microbiome Project data focusing on Bacteroides and Faecalibacterium genera.

**Figure 3.**
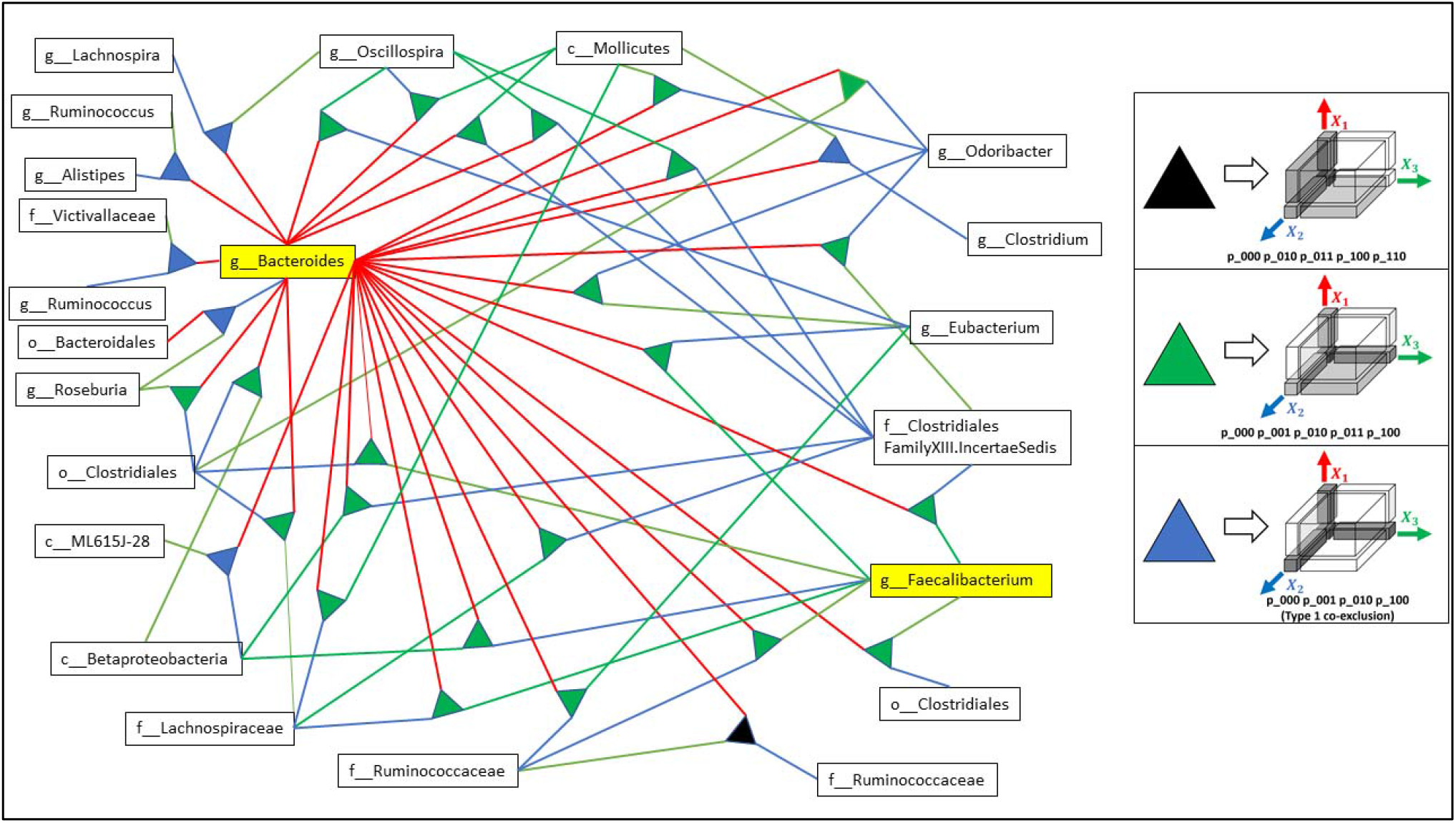
3D patterns in the Human Microbiome Project stool data focusing on Bacteroides and Faecalibacterium genus.

### 3D Patterns in *H. pylori* Gene Composition Associated with Duodenal ulcers, Gastritis, and Gastric ulcers

Fully annotated *Helicobacter pylori* genomes (93 total) were downloaded from the PATRIC BRC database ^7^. The database included 58 *H. pylori* genomes from chronic gastritis, 21 from gastric ulcers, and 13 from duodenal ulcers. The features of every genome were extracted and merged based on their protein product annotation from Patric BRC database. The unannotated features were excluded from the analysis as they could not be merged across samples based on the annotation identifier.

The focus of the analysis was to identify only patterns that include diagnosis type metadata, so, in order to be considered, patterns rewire to include duodenal ulcer, gastritis, or gastric ulcers as one of the features. While no 2D patterns were identified with a minimal pattern score of 0.99 and a population threshold of 0.1, total of 28 3D patterns (Supplementary Table 3 and Figure 4) have been detected in the dataset.

**Figure 4.**
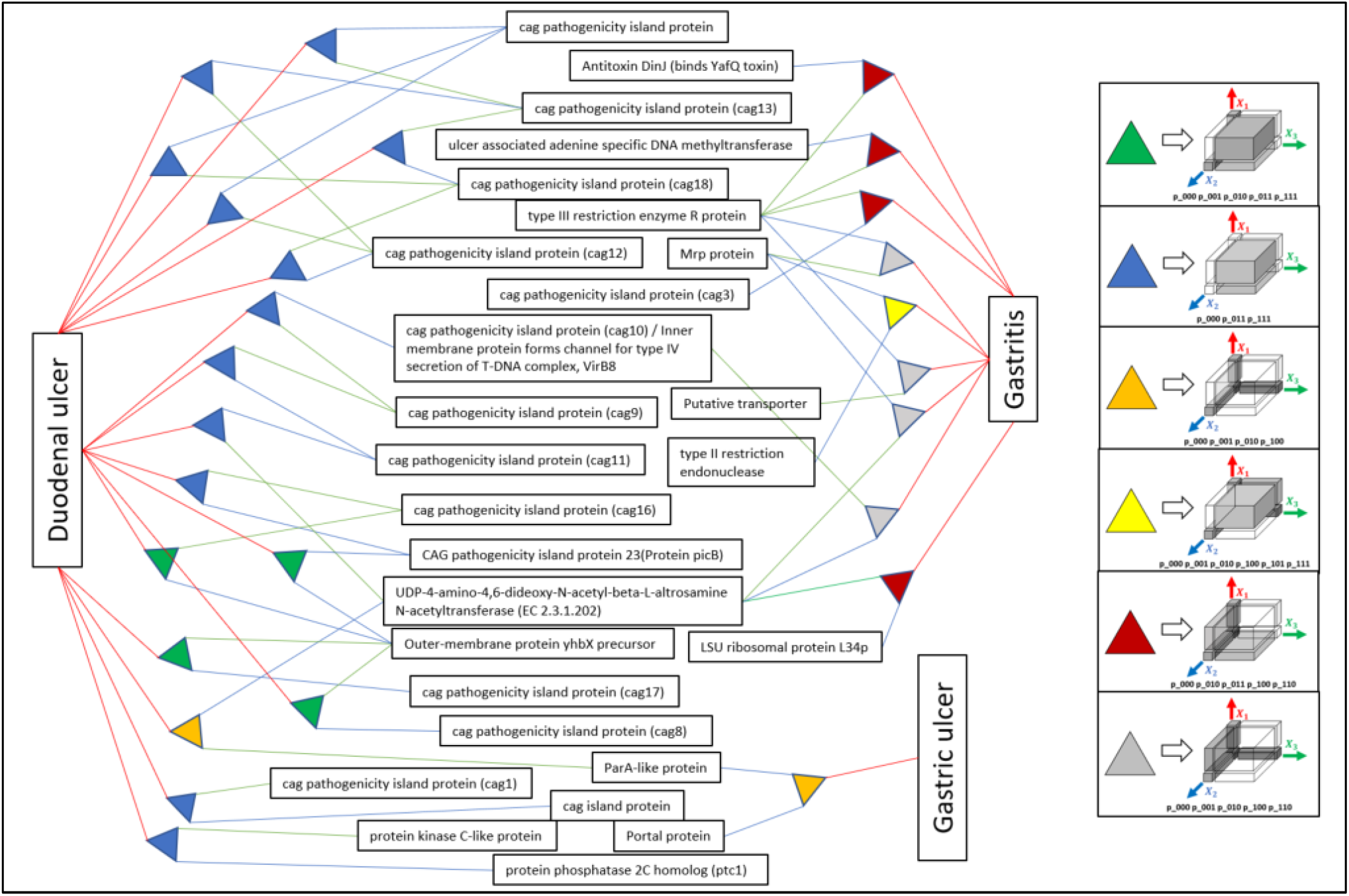
Subset of 3D patterns in *H. pylori* gene composition associated to duodenal ulcers, gastritis, and gastric ulcers.

## Discussion

In contrast with dimensionality reduction (such as principal components analysis and multidimensional covariation) and AI (neural network, deep learning, etc.) solutions, patterns identified by the proposed approach may provide better background for meaningful mechanistic interpretation of the biological processes. Additionally a variety of mutual information-based methods are not suitable for estimating the strength of Boolean patterns because of the effects of the number of populated partitions and disbalance of the partitions’ population on the pattern’s score ^1^. Our preliminary analysis suggests that multidimensional patterns may not just be present but could dominate the landscape of multi-omics data, which is not surprising because complex interactions between components of biological systems (microbiomes, gene regulations, etc.) are unlikely to be reduced to simple pairwise interactions.

We also believe that different patterns represent different properties of biological systems: while some patterns, such as *co-presence* or *one-way relations* can be interpreted as an interaction between system's components (proteins, microorganisms, etc.); other patterns, such as *co-exclusion* may represent the "design rules" of the systems: for example allow predicting which microbiome can and cannot exist. Interestingly enough (data not shown), an artificial dataset can be constructed to simultaneously fit with the highest possible (100%) score to more than one pattern. If detected in the real data, such an observation may reflect different activities in which these features can be involved.

It is also necessary to mention that the proposed approach can be extended to more complex discrete patterns by using 3, 4, etc. thresholds for each feature simultaneously, so instead of absence/presence thresholds, one can consider patterns containing several levels of abundance, such as absence and presence in low, medium, and high abundance. In which case will optimization of 3 thresholds: *ε*_*absence*_; *ε*_*low*_; *ε*_*medium*_; for each feature would be necessary.

It is important to keep in mind that search for multidimensional patterns (as a search for any other type of patterns) is exploratory. Each finding must be validated. The most important outcome of the proposed analysis is that many hypotheses are filtered out by this approach so one can focus validation effort on a much smaller number of patterns.

